# The Molecular Signature of Umami Palatability in Dogs Based on Amino Acid Interactions with Canine Taste Receptors

**DOI:** 10.1101/2025.10.13.681969

**Authors:** Margaux Brewer, Arun HS Kumar

## Abstract

Canine feeding behaviour is strongly influenced by taste perception, yet the molecular determinants of palatability, particularly umami taste is unclear. Dogs typically consume food rapidly with minimal mastication, relying on a limited number of taste buds to detect sour, bitter, salty, sweet, and umami flavours. Amino acids are known to play a central role in canine taste, especially in diets rich in animal proteins. In this study, we aimed to identify the specific amino acids that optimally stimulate umami perception in dogs using a receptor–ligand docking approach. A total of 27 canine-specific taste receptors were identified from the UniProt database, comprising three umami, two sweet, five uncharacterized, and seventeen bitter receptors. All 20 naturally occurring amino acids were docked against these receptors using the CB-Dock tool, and binding affinities were systematically analysed. Heat maps of binding energies revealed that tyrosine, tryptophan, arginine, histidine, phenylalanine, glutamine, glutamic acid, and lysine exhibited the strongest interactions with umami and sweet receptors, whereas bitter and unknown receptors showed comparatively weaker affinities. Binding energy ratio analyses further confirmed that amino acids preferentially stimulated umami and sweet receptors, with lysine, histidine, glutamine, glutamic acid, and arginine emerging as key co-stimulators. Functional enrichment analyses indicated that the receptors belong to Class C/3 GPCRs, with membrane-bound sweet receptor complexes and strong associations with sensory perception of umami, sweet, and bitter tastes. These findings provide a molecular basis for enhancing palatability in canine diets, offering practical implications for the formulation of dry dog food and the design of oral veterinary medicines. The study highlights the importance of specific amino acids, potentially in combination with salts, in modulating taste responses and improving food acceptance in dogs.

## Introduction

Domestic dogs exhibit unique feeding behaviours characterized by rapid consumption of food with minimal mastication or savouring.^1,2^ This gluttonous eating pattern is thought to be associated with their relatively limited gustatory capacity; dogs possess only a fraction of the taste buds found in humans.^3,4^ Despite this, dogs can detect the primary taste modalities (sour, bitter, salty, sweet, and umami) when the corresponding chemoreceptors are activated.^2,5^ Among these, umami perception is of particular interest, as it is closely linked to the detection of amino acids and nucleotides, components abundant in animal tissues and central to canine dietary preferences.

Food selection in dogs is influenced not only by nutritional adequacy but also by short-term palatability. Palatability is shaped by an interplay of sensory cues, including taste, aroma, texture, size, appearance, temperature, and consistency.^6-8^ In addition, canine food preferences can be modulated by genetics and early-life experiences, further complicating efforts to optimize commercial formulations.^8^ Importantly, dogs exhibit heightened sensitivity to the taste of amino acids, organic acids, and nucleotides, suggesting a key role of these molecules in palatability-driven feeding behaviour.^5,9^ From a product development perspective, managing palatability in dry pet foods presents unique challenges. Such diets typically consist of carbohydrate sources (e.g., cereals), protein sources (fresh, frozen, or rendered meals derived from animal tissues), by-products, and lipid sources of animal or plant origin.^3^ The nature of these ingredients, along with their processing and incorporation methods (such as blending, concentration, or surface coating), critically influences overall product appeal.

Commercially popular and highly rated dry dog foods utilize a diverse range of protein ingredients. Animal-derived proteins such as chicken, turkey, salmon, fish, and mixed poultry are staples in many dry formulations, while beef, lamb, venison, duck, and rabbit are also frequently incorporated to broaden flavour profiles and cater to varying consumer demands.^10-13^ More recently, novel and nontraditional protein sources have emerged, particularly in diets marketed for dogs with sensitivities or requiring alternative protein options.^11,13,14^ The choice of protein not only ensures essential amino acid provision but also directly affects palatability, as different animal tissues contribute distinct sensory characteristics. In addition to protein type, the molecular composition of diets plays a critical role in influencing canine taste perception.^3^ Specific amino acids including L-phenylalanine, L-tyrosine, L-tryptophan, L-methionine, L-arginine, L-leucine, and L-serine are particularly important for enhancing flavour responses and improving palatability.^5,15^ Furthermore, salts such as sodium, potassium, and calcium can synergistically intensify the taste responses elicited by these amino acids, thereby amplifying overall product appeal.^16,17^ By incorporating these key factors into dry food formulations, manufacturers aim to achieve a balance between nutritional adequacy and sensory acceptance, which remains a central challenge in the design of canine diets.

Although a wide range of meat sources are used in the formulation of commercial canine diets to diversify flavour and nutritional value, the underlying mechanisms that govern palatability at the molecular level remain poorly understood.^10,11^ Current formulations often rely on empirical ingredient selection rather than a systematic evaluation of how specific amino acids interact with canine taste receptors to drive umami perception. Despite the central role of amino acids in enhancing flavour, a receptor–ligand-based framework for identifying which amino acids most effectively stimulate umami taste in dogs has not yet been developed. Such an approach would provide valuable insights into the molecular drivers of taste perception and enable the rational design of canine diets with optimized palatability. Given the centrality of amino acid perception in canine feeding, the present study focused on systematically evaluating the interactions between all 20 naturally occurring amino acids and the complete repertoire of known canine taste receptors through molecular docking approaches. This framework provides a mechanistic basis for understanding the molecular determinants of umami-driven palatability in dogs and offers insights for the rational use of palatability factors in dry dog foods to enhance acceptance.

## Methods

All dog-specific taste receptors reported in the literature and available UniProt databases were included in this study. The initial step involved the compilation of the complete repertoire of canine taste receptors, including those associated with sweet, sour, salty, bitter, and umami perception. Once collated, the receptor genes were subjected to protein–protein interaction (PPI) network analysis using STRING (Search Tool for the Retrieval of Interacting Genes/Proteins, version 12.0). STRING was selected due to its robust capacity to predict both direct (physical) and indirect (functional) associations, integrating information from curated databases, experimental data, co-expression, and text mining. To ensure high confidence in the results, interaction scores were filtered using a minimum confidence threshold of 0.7, and only experimentally validated or strongly predicted interactions were retained for subsequent analyses. The resulting PPI networks were then visualized and topologically analysed to identify hub nodes, clusters, and modules that could represent key signalling pathways involved in taste perception. Network statistics such as degree centrality, betweenness centrality, and clustering coefficients were computed to assess the relative importance of specific receptors within the network. To obtain biological insights from the identified receptor interaction networks, gene ontology (GO) enrichment analysis was performed. The analysis was conducted using the GO functional annotation tools integrated within STRING (version 12.0) for cross-validation. Enrichment was assessed across three GO categories: biological processes, molecular functions, and cellular components. Statistical significance of enriched GO terms was determined using the Benjamini–Hochberg false discovery rate (FDR) correction, with an adjusted p-value threshold of <0.05 considered significant. The integrated approach of network analysis and GO enrichment provided a systems-level understanding of the functional landscape of canine taste receptors, laying the foundation for further receptor–ligand docking studies to identify amino acids capable of optimally enhancing umami palatability in dogs.

A total of 27 canine-specific taste receptors were identified and retrieved from the UniProt database. These receptors included members of the bitter (TAS2R), sweet/umami (TAS1R2/TAS1R3,1), and other chemosensory receptor families known to be functionally relevant in dogs. Protein sequences were obtained in FASTA format, and corresponding three-dimensional (3D) structures were modelled where experimental crystal structures of full-length protein were not available as reported before.^18,19^ Homology modelling was performed using SWISS-MODEL, and the generated models were validated based on structural quality assessment parameters such as GMQE, QMEAN, and Ramachandran plot statistics. Where available, experimentally resolved structures from the Protein Data Bank (PDB) were used directly.

Of the 27 receptors identified, three were classified as umami-associated, two as sweet-associated, five were categorized as orphan or unknown receptors with uncharacterized specificity, and the remaining 17 represented bitter taste receptors. This distribution highlights the evolutionary emphasis on bitter taste perception in dogs, likely as a protective mechanism against the ingestion of harmful compounds, while still retaining a smaller but functionally critical set of receptors for detecting amino acid and nucleotide-based flavours such as umami.

All 20 naturally occurring amino acids were considered as ligands for docking studies. The 3D structures of the amino acids were retrieved in SDF format from the PubChem database and converted to PDB file and energy-minimized using Chimera (version 1.19) to ensure geometrical stability prior to docking.^18-20^ Molecular docking was carried out using the CB-Dock tool, an automated web server designed for cavity detection-guided blind docking. Each receptor was subjected to cavity detection to identify potential ligand-binding sites, after which docking simulations were performed for all 20 amino acids against each receptor. CB-Dock employs AutoDock Vina as its docking engine, which calculates binding affinities based on predicted free binding energies (kcal/mol).^18-20^ For each receptor–ligand pair, the top-ranked docking poses were selected based on binding energy scores. The docking results were systematically compiled to identify the amino acids with the strongest predicted binding affinities across the canine taste receptor repertoire. Emphasis was placed on the receptors associated with umami taste (e.g., TAS1R1 and TAS1R3), though comparisons were made across all receptor families to assess broader selectivity trends. The binding interactions of the top amino acids were visualized and analysed using Chimera (version 1.19) to examine hydrogen bonding, hydrophobic interactions, and other non-covalent interactions stabilizing the receptor–ligand complexes.^18,19^ The docking simulations produced binding energy values (kcal/mol) for each receptor–amino acid pair, representing the predicted affinity of the ligand toward the respective canine taste receptor. These values were systematically compiled into a receptor–ligand interaction matrix, where column represented the 27 taste receptors and row represented the 20 naturally occurring amino acids. The compiled dataset allowed for comparative evaluation of binding affinities across the receptor repertoire.

To facilitate interpretation, the binding energy data were visualized as heat maps. Heat map construction was performed using the R statistical environment (version 4.42) with the “pheatmap” package, which enabled hierarchical clustering of both receptors and ligands based on their affinity profiles.^18,19^ Colour gradients were applied, with lower (more negative in green colour) binding energy values indicating stronger affinities, thereby visually highlighting amino acids with preferential binding to specific receptors. Clustering analysis was used to identify groups of amino acids sharing similar binding patterns as well as receptor subgroups with overlapping affinity profiles. This heat map approach provided a clear comparative framework to assess the relative affinities of amino acids across all identified canine taste receptors, allowing for the recognition of key ligand–receptor interactions potentially associated with umami taste enhancement.

To further investigate the selective preference of specific amino acids for umami taste receptors, binding energy ratio analyses were performed. For each amino acid, the receptor with the highest binding affinity (lowest binding energy) was identified and used as the normalization reference. Binding energy values of the same amino acid across all other receptors were then expressed as ratios relative to this reference value. This normalization allowed for the assessment of amino acid selectivity by controlling for inherent differences in docking scores. The normalized binding energy ratios were subsequently grouped according to receptor type category, namely umami, sweet, bitter, and unknown receptors. For each category, the average ratio was calculated, enabling comparison of how strongly individual amino acids were predicted to interact with umami receptors relative to other receptor classes. This analysis provided a quantitative framework to identify amino acids most likely to contribute to strong umami-driven taste perception in dogs.

As an additional layer of assessment, pairwise binding energy ratios were computed between receptor categories to evaluate the dominant taste sensation potentially triggered by each amino acid. Specifically, the following category pairs were examined: umami/sweet, umami/bitter, umami/other, sweet/bitter, sweet/other, and bitter/other, (where “other” denoted the group of five unknown receptors). These ratio comparisons highlighted the relative strength of amino acid interactions across different taste modalities, offering insight into whether a given amino acid preferentially activates umami-associated receptors versus alternative taste receptor classes. This ratio-based approach allowed for a more nuanced evaluation of receptor–ligand specificity beyond raw binding energy values, facilitating the identification of amino acids most likely to enhance umami palatability in canine diets.

## Results

The network analysis of canine taste receptors revealed a highly interconnected protein–protein interaction (PPI) network. The constructed network comprised 15 nodes and 83 edges, with an average node degree of 11.1, indicating that each receptor was connected to multiple partners within the group. The average local clustering coefficient was 0.899, suggesting a high tendency for receptors to form tightly clustered interaction neighbourhoods rather than existing as isolated units. The analysis further showed an expected number of edges of 0, while the observed number of edges was substantially higher. The PPI enrichment p-value was < 1.0e-16, confirming that the observed interactions were highly significant and not due to random chance. This enrichment strongly indicates that the canine taste receptor proteins share functional and biological relationships, forming a cohesive interaction network. Collectively, these findings suggest that canine taste receptors are not independent entities but rather participate in a biologically meaningful, interconnected system, potentially underlying coordinated roles in taste perception and related signalling pathways. We also observed a dense subnetwork formed by the TAS1R family suggesting the strong integration of sweet and umami pathways, reflecting attraction to both sugars and protein-rich foods in dogs. In contrast, the bitter receptor cluster are positioned more peripherally, consistent with a carnivorous dietary bias and diminished need for broad detection of plant-derived toxins. The network architecture illustrates an evolutionary balance in canine taste perception, preserving attraction to calorie- and protein-dense foods.

Tissue enrichment analysis indicated that the identified canine taste receptors were exclusively expressed in taste buds, with the majority classified as type 2 taste receptors. Receptor Classification and Tissue Mapping (RCTM) analysis demonstrated high enrichment for sensory perception of sweet, umami (glutamate) and bitter, taste modalities, consistent with their classification as Class C/3 (metabotropic glutamate/pheromone receptors) with G alpha (i) signalling pathways. Further receptor annotation confirmed that all identified canine taste receptors belong to the Class C/3 GPCR, family 3 group. Functional enrichment data suggested that both general taste receptor activity and bitter taste receptor activity were equally prominent, highlighting the evolutionary role of bitter perception in dogs alongside nutrient-driven modalities. Compartment enrichment analysis further indicated receptor association with membrane-bound sweet taste receptor complexes, suggesting structural relevance for receptor clustering and signalling efficiency.

The docking analysis of 20 naturally occurring amino acids against the 27 canine taste receptors revealed several key trends. Tyrosine, tryptophan, arginine, and phenylalanine consistently exhibited the highest binding affinities across most taste receptors, underscoring their broad role in canine taste perception. When examined by receptor type, tyrosine, tryptophan, arginine, histidine, phenylalanine, glutamine, glutamic acid, and aspartic acid demonstrated strong binding specifically to umami-associated receptors. In contrast, tyrosine, tryptophan, phenylalanine, isoleucine, leucine, lysine, histidine, glutamine, glutamic acid, and arginine displayed high affinity toward sweet receptors. For bitter and unclassified (orphan) receptors, tyrosine, tryptophan, and phenylalanine were identified to have strongest affinities. These results aligned with prior reports identifying tyrosine, tryptophan, serine, methionine, leucine, and arginine as major stimulators of umami taste receptors, thereby validating the docking outcomes.

To further delineate amino acid selectivity, binding energy ratios were calculated by normalizing each amino acid’s receptor affinities against its strongest binding interaction. This analysis revealed that both sweet and umami receptor categories were equally and strongly stimulated by amino acids, with significantly higher ratios than those observed for bitter and unknown receptor types. The average binding energy across receptor categories supported this pattern, with lysine, histidine, glutamine, glutamic acid, and arginine emerging as major co-stimulators of both sweet and umami receptors. Pairwise ratio comparisons between receptor categories provided additional insight into the relative dominance of taste responses. The umami/bitter, umami/other, sweet/bitter, and sweet/other groups exhibited significantly higher ratios, indicating that amino acids preferentially stimulated sweet and umami receptors over bitter and unknown receptors. Notably, the umami/sweet ratio was close to 1.0, suggesting equal stimulation of both receptor types by a broad range of amino acids. The unknown receptor group consistently showed the weakest binding affinities and lowest ratio values, reinforcing their limited role in amino acid-mediated taste perception. Collectively, this relative affinity framework identified lysine, histidine, glutamine, and glutamic acid as the most potent and selective co-stimulators of umami and sweet receptor classes in dogs.

## Discussion

This study provides a comprehensive receptor–ligand-based framework to investigate the molecular basis of umami and sweet taste perception in dogs through docking of 20 naturally occurring amino acids against 27 canine-specific taste receptors. Our findings demonstrated that amino acids such as lysine, histidine, glutamine, and glutamic acid play prominent roles in selectively stimulating umami and sweet receptors, while also exhibiting comparatively weaker affinities toward bitter and unknown receptor classes. These results highlight the dual importance of umami and sweet perception in shaping canine food preferences and provide a molecular foundation for improving palatability in commercial diets. These findings carry important implications for canine nutrition and veterinary medicine. Pet dogs are often exclusively or partially fed diets consisting of commercially prepared foods, where palatability is a key determinant of acceptance.^21,22^ As the pet food industry shifts toward sustainable protein sources, including alternative and non-animal proteins, palatability challenges are likely to increase.^23,24^ The identification of amino acids that selectively stimulate sweet and umami receptors provides a molecular basis for enhancing acceptance of such formulations. Furthermore, the strong aversion of dogs to bitter tastes poses a challenge in oral delivery of veterinary medicines.^25,26^ Understanding the receptor-level drivers of taste perception could inform the design of formulations that mask bitterness while leveraging amino acid-mediated stimulation of sweet and umami receptors to improve compliance in oral drug administration.

The network and enrichment analyses provide new insights into the organization and functional relevance of canine taste receptors. The highly interconnected PPI network observed in this study, with significant enrichment well beyond random expectations, indicates that taste receptors in dogs operate as a coordinated system rather than isolated sensory units. This observation is consistent with previous findings in humans and rodents, where taste receptors have been shown to interact within broader signalling frameworks to fine-tune perception.^27,28^ The dense subnetwork formed by TAS1R family members suggests a strong functional integration of sweet and umami pathways. This may reflect the dual importance of sugars and amino acid–rich foods in canine diets, consistent with evidence that dogs are particularly sensitive to amino acids, nucleotides, and other compounds abundant in animal tissues.^3^ The peripheral placement of bitter receptors in the network, by contrast, aligns with the carnivorous origins of dogs, where the selective pressure for broad detection of plant-derived toxins is reduced compared to omnivores. This positioning suggests that bitter receptors, while still functionally relevant, play a secondary role relative to umami and sweet perception. The enrichment analyses further support this interpretation by showing exclusive expression of receptors in taste buds, high representation of Class C/3 GPCRs, and strong associations with sweet, umami, and bitter modalities. Notably, the equal prominence of general and bitter taste receptor activities echoes prior work highlighting the evolutionary conservation of bitter perception as a safeguard, even in carnivorous species.^29-31^ The evolution of canine taste receptors as a tightly integrated network that prioritizes attraction to nutrient-rich foods while maintaining the ability to detect aversive compounds offers a critical insights into development of highly palatable diet.

Our observation of a strong functional integration between the sweet and umami pathways in the network analysis aligns with the finding that both receptor classes were equally stimulated by amino acids. This relationship further correlates with the physiological feeding behaviour of dogs.^2,32^ Unlike humans, dogs do not invest much time in mastication or savouring but instead consume food rapidly and in a gluttonous manner.^33,34^ This behavioural tendency is accompanied by a reduced number of taste buds compared to humans.^3^ Despite this limitation, dogs can detect all five primary taste modalities, and the strong stimulation of sweet and umami receptors may compensate for their limited gustatory capacity, ensuring efficient detection of nutrient-rich foods.^35^ Given that amino acids, organic acids, and nucleotides are abundant in animal tissues, the heightened canine sensitivity to these compounds is consistent with an evolutionary adaptation toward a carnivorous-leaning diet.^9^ The implications of our findings extend directly to the formulation of commercial dry dog food. Palatability in dogs is not determined solely by taste, but by a combination of factors including aroma, texture, size, appearance, temperature, and consistency.^3^ Our study emphasizes that amino acids with strong selective umami and sweet receptor stimulation, particularly lysine, histidine, glutamine, and glutamic acid could serve as key molecular drivers of enhanced food acceptance. Several amino acids (tyrosine, tryptophan, arginine, and phenylalanine) have been consistently identified as key contributors to palatability in canine diets.^5,9^ The importance of these amino acids has been linked to their strong sensory impact and prevalence in animal-derived protein sources, which dominate most commercial formulations. Interestingly, while our results confirmed the strong binding potential of these amino acids, their interactions were not restricted to a single taste modality. Instead, tyrosine, tryptophan, arginine, and phenylalanine demonstrated comparable affinities across sweet, umami, bitter and unknown receptor classes. This broad responsiveness suggests that these amino acids may function as general taste modulators, contributing to multiple sensory pathways rather than selectively driving umami or sweet perception. However, their non-selective affinity towards bitter receptors also raises an important consideration i.e., foods rich in tyrosine, tryptophan, arginine, and phenylalanine may not always translate into truly palatable diets for dogs. While these amino acids can stimulate nutrient-positive tastes such as sweet and umami, their concurrent activation of bitter receptors may introduce aversive signals that diminish overall acceptance. Bitter taste perception in dogs, as in many mammals, is often associated with the detection of potentially harmful or toxic compounds.^29,36^ Therefore, the dual stimulation of both positive (umami/sweet) and negative (bitter) pathways may result in a mixed sensory experience, limiting the net palatability of diets enriched with these amino acids. This finding contrasts with the more selective interactions observed for lysine, histidine, glutamine, and glutamic acid, which preferentially target sweet and umami receptors while avoiding strong activation of bitter receptors. Such selective receptor engagement provides a clearer molecular rationale for why these amino acids may be more effective drivers of palatability in canine diets, offering a taste profile that reinforces positive sensory cues without simultaneously triggering aversion. The distinction between broadly binding amino acids and those with selective receptor preferences is an important finding in our study, as it provides a mechanistic explanation for why certain amino acids can effectively contribute to acceptance of dog food formulations. Moreover, the selective activation of umami and sweet receptors by lysine, histidine, glutamine, and glutamic acid may offer a more targeted strategy for enhancing palatability in future diet design, especially in the context of alternative protein sources or oral drug formulations where masking bitterness is critical. From a nutritional and product development standpoint, these insights are particularly important as the pet food industry increasingly incorporates alternative and non-animal protein sources to improve sustainability. While such proteins may offer environmental benefits, they may lack the same amino acid profiles as traditional animal-derived sources, leading to reduced palatability. Our understanding of which amino acids most effectively stimulate umami and sweet receptors, provides a rational basis for designing palatability enhancers tailored for canine diets.

## Figure legends

**Figure 1:**
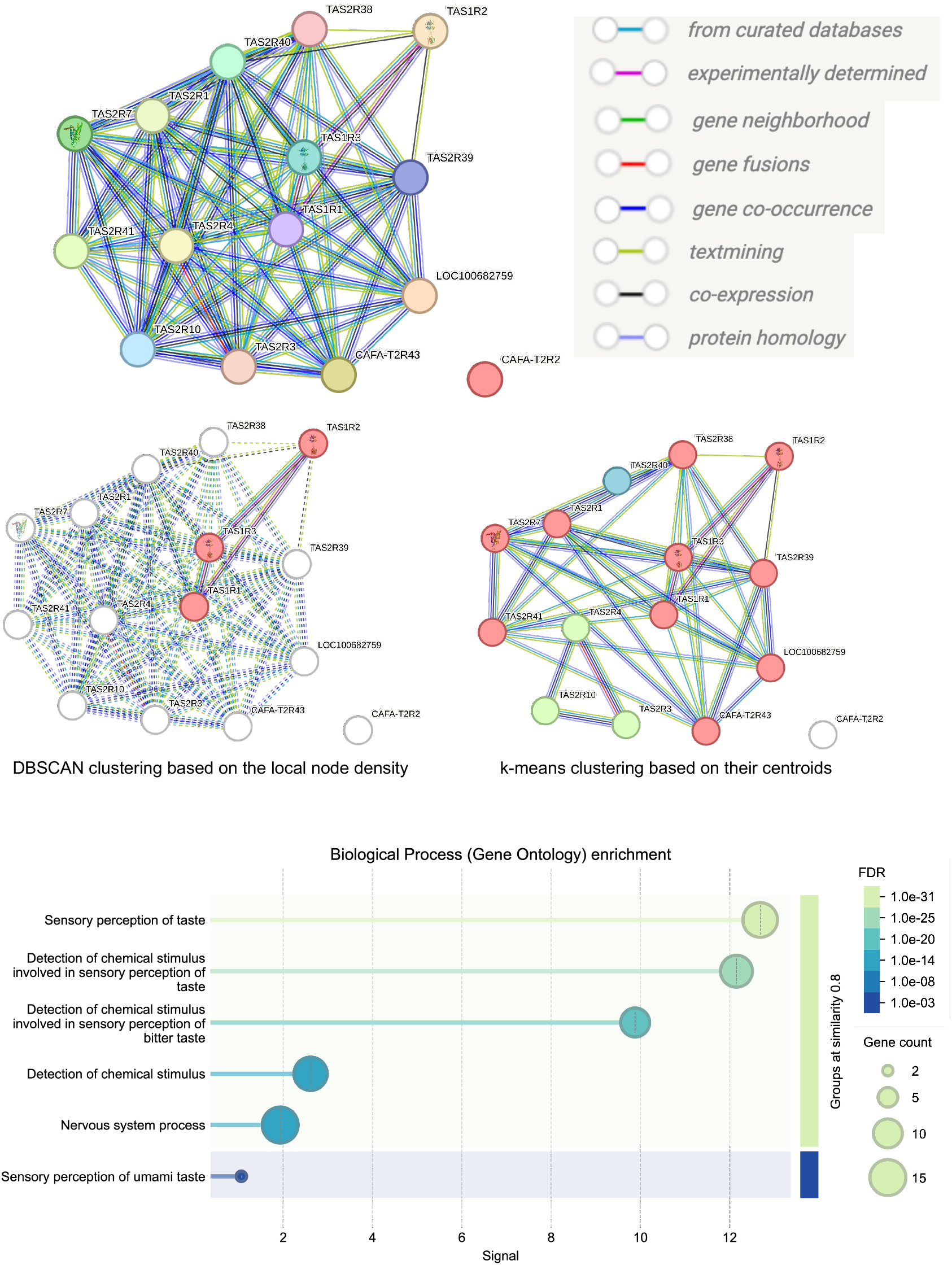
Network analysis and functional enrichment of canine taste receptors. Each node represents an individual taste receptor encoded by a single, protein-coding gene locus with edges indicating predicted or experimentally validated protein–protein interactions. Canine taste receptors were further organized based on local node density and centroid-based clustering to highlight functional subnetwork relationships. The bubble plot represents Gene Ontology (GO) enrichment analysis of the network, displaying significantly enriched biological processes. False discovery rate (FDR).

**Figure 2:**
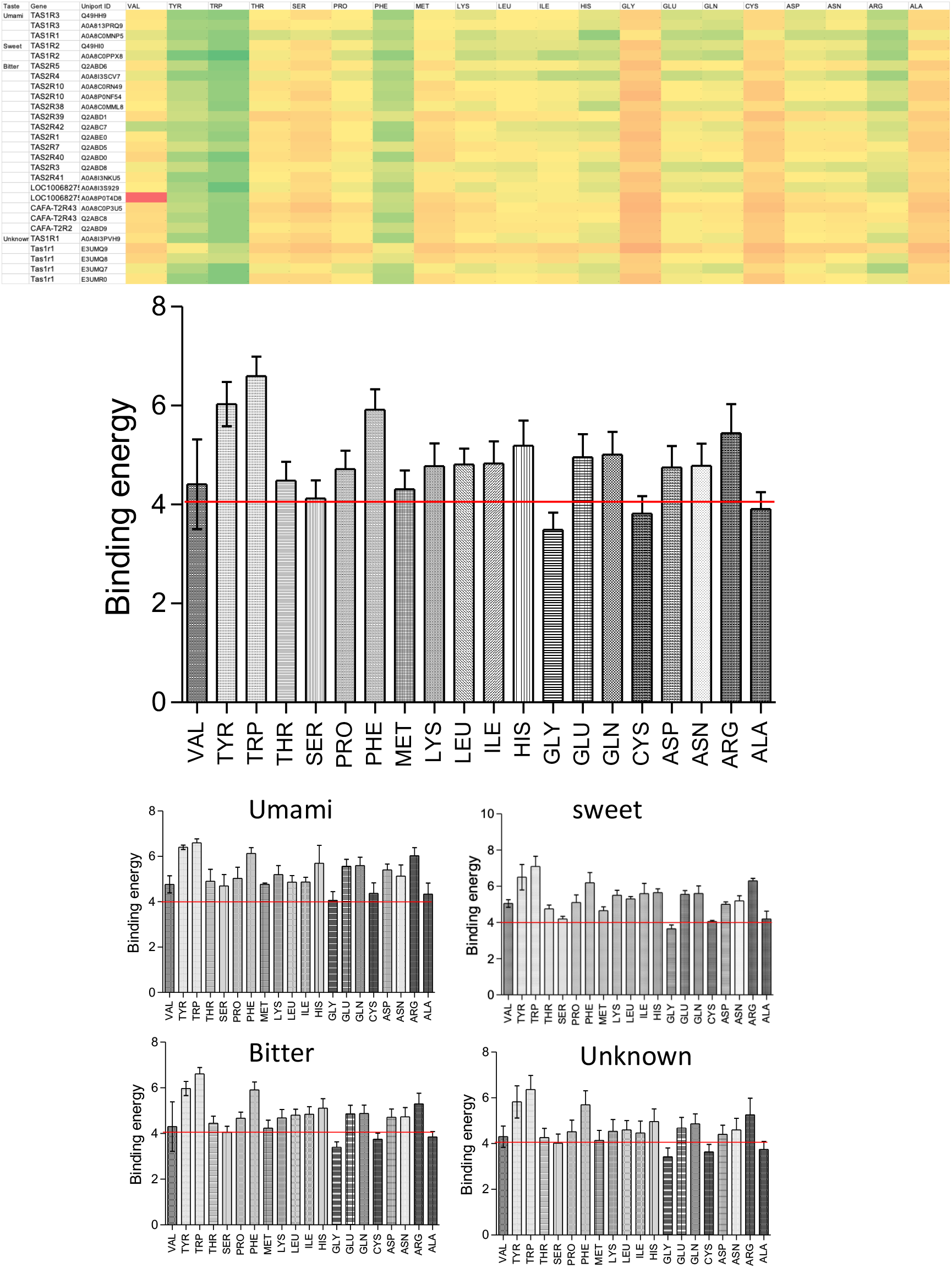
Heatmap and comparative binding energy profiles of amino acids across canine taste receptor classes. The heatmap illustrates the binding energy values of all 20 naturally occurring amino acids docked against the full set of canine taste receptors, highlighting differential affinities across receptor types. Each cell represents the predicted binding energy (kcal/mol), where lower values (green) indicate stronger ligand–receptor interactions. The bar graphs display the average binding energy (mean ± SD) of each amino acid with all canine taste receptors combined, as well as separately for umami, sweet, bitter, and unknown receptor categories.

**Figure 3:**
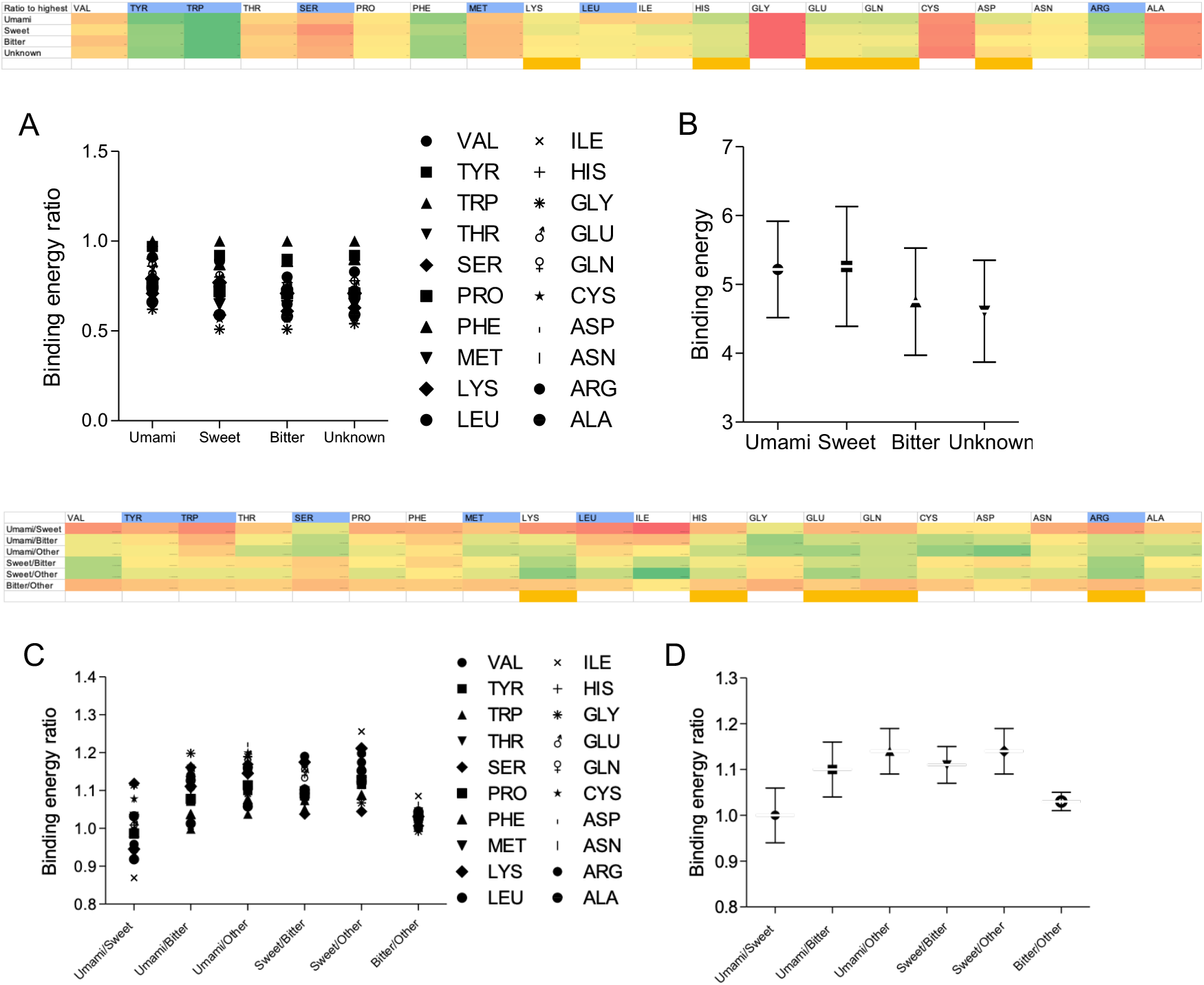
Comparative analysis of amino acid binding energy ratios across canine taste receptor categories. The heatmap illustrates the binding energy ratios of 20 naturally occurring amino acids with the four major classes of canine taste receptors: umami, sweet, bitter, and unknown. Corresponding bar graphs display (A) the mean binding energy ratios and (B) the average binding energy values (kcal/mol) of each amino acid across receptor categories, presented as mean ± SD. The lower heatmap depicts the binding energy ratios of the same 20 amino acids across pairwise receptor comparisons (umami/sweet, umami/bitter, umami/unknown, sweet/bitter, sweet/unknown, and bitter/unknown). Bar graphs show (C) the mean binding energy ratios and (D) the average of these ratios (mean ± SD) for each amino acid pairwise comparison.

